# A spectral framework for measuring diversity in multiple sequence alignments

**DOI:** 10.64898/2026.02.09.704934

**Authors:** Vaitea Opuu

## Abstract

Machine learning (ML) methods for proteins and RNAs rely on multiple sequence alignments (MSAs) and related datasets such as experimental mutagenesis libraries, yet the amount of usable information they contain remains unclear. Here, a spectral measure of information is recast into an interpretable quantity for MSAs, denoted *L*_*eff*_, defined as the number of fully independent alignment positions that reproduce the observed sequence diversity. Applied to RNA MSAs, this measure shows that evolutionary constraints nearly halve diversity relative to the secondary structure alone, quantifying functional and phylogenetic restrictions beyond base pairing. The same analysis indicates even lower effective diversity in proteins, quantifying stronger physicochemical and evolutionary constraints on amino acids. *L*_*eff*_ further correlates with protein structure prediction accuracy, anticipating cases with insufficient evolutionary signal. When applied to experimentally and computationally generated libraries, it measures both produced diversity and cross-library overlap, quantifying novelty rather than redundant sampling. Together, these results establish *L*_*eff*_ as an operational tool to estimate effective information in MSAs, anticipate modeling difficulties, and guide protein and RNA design.

## Introduction

Modern sequencing and design technologies now generate massive collections of biological sequences [1], especially for RNA [2]. These data are the primary input for machine learning (ML) models used to infer evolutionary constraints or generate functional variants. Such models support applications including mutational effect prediction from deep mutational scanning [3] and protein structure prediction from natural multiple sequence alignments (MSA), as in AlphaFold [4]. Model performance is often assumed to scale with dataset size [5], yet the amount of usable information contained in these data is usually unknown. Natural MSAs are shaped by phylogeny and uneven sampling, resulting in many closely related sequences [6, 7]. Designed or experimental libraries, in contrast, densely explore narrow mutational neighborhoods around a reference [2]. Consequently, large datasets can have limited effective diversity. Estimating the information content is therefore necessary to set expectations for model performance, to assess whether new data add genuinely novel information, and to relate model capacity to the diversity present in the training data.

Several measures are commonly used to summarize sequence diversity, but their interpretation and relevance for model behavior are often unclear. Average pairwise (Hamming) distance measures how different two randomly chosen sequences are on average but can be biased by sequence clusters. Positional Shannon entropy quantifies variability at individual alignment positions [8], which is informative about local conservation but treats positions independently. Similarly, k-mer entropy summarizes the diversity of short subsequences [9], but remains insensitive to long-range constraints. The effective number of sequences downweights clusters of nearly identical sequences to correct for redundancy [10], yet depends on an arbitrary similarity threshold and yields a dataset size whose meaning for model performance is not clear [11]. As a result, these measures often yield different views of diversity because they probe only local patterns or partial summaries of variation. An ideal measure of diversity would use the MSA as a sample to estimate the effective diversity of the underlying sequence distribution, rather than merely summarizing observed variation.

In this work, I introduce an information-based measure of sequence diversity that treats an MSA as a sample and provides a dataset-level estimate of the effective information contained in the underlying sequence distribution. The resulting quantity, the effective sequence length *L*_*eff*_, is designed to capture global variation across the full alignment rather than local patterns or raw sequence counts. By construction, *L*_*eff*_ depends on the collective structure of variation across sites and thus reflects correlations and constraints distributed along the sequence. Interpreting diversity on an effective-length scale allows different MSAs and experimental libraries to be compared, providing an intuitive and model-derived summary of their overall information content.

The paper is organized as follows. I first review existing measures of sequence diversity in aligned data. I then introduce the effective sequence length *L*_*eff*_ and describe its properties. Next, I analyze its behavior on synthetic datasets with controlled constraints to show that it captures global variation induced by correlations in the MSA. I then apply the measure to natural RNA and protein MSAs and to experimentally designed ribozyme libraries, highlighting differences in how variation is distributed across sequence space. The implementation and data are available at https://github.com/vaiteaopuu/effective_length.

## Available diversity measures

Diversity and information in an MSA are not uniquely defined quantities, but rather a collection of nonequivalent summaries of variability in sequence space. Existing measures differ in the scale at which they operate (site-wise, sequence-wise, pairwise, or local contexts), in whether they rely on an explicit statistical model or are purely empirical, in their dependence on user-defined parameters, and in the type of structure they are sensitive to.

### Positional entropy

Positional entropy quantifies sequence variability at the level of individual alignment positions. Let *p*_*ia*_ denote the empirical frequency of symbol *a* at position *i*. The positional Shannon entropy is defined as

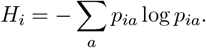

Under an independent-site assumption, these positional entropies can be combined into a global support size,

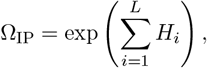

which corresponds to the number of distinct sequences expected. These measures are effectively parameter-free aside from alphabet definition and gap handling, and are computationally inexpensive. They are widely used to assess positional conservation in MSAs and, more generally, as measures of uncertainty or support size in probabilistic sequence models, including large language models. Their main limitation is that they neglect inter-site correlations, typically leading to an overestimation of the effective information content.

### *k*-mer diversity

*k*-mer diversity measures variability at the level of short, local sequence patterns. For all overlapping subsequences of length *k*, let *q*_*m*_ denote the empirical frequency of each distinct k-mer (m) across the MSA. The Shannon entropy

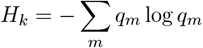

defines an effective number of k-mers as *exp*(*H*_*k*_). This measure is explicitly parameterized by the choice of *k* and is commonly used to quantify motif richness and local sequence complexity in MSAs. While easy to compute for small *k*, the exponential growth of the *k*-mer alphabet leads to sparsity and estimation issues as *k* increases. As with positional entropy, this measure is restricted to local patterns and does not capture long-range dependencies.

### Effective number of sequences

The effective number (*N*_*eff*_) of sequences measures diversity correcting for redundancy in an MSA. Let *s*_*ij*_ denote the fractional identity between sequences *i* and *j*, and define neighbors as sequence pairs whose similarity exceeds a fixed threshold, typically set to 80% [10]. Each sequence *i* is assigned a weight

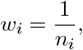

where *n*_*i*_ is the number of its neighbors. The effective number of sequences is then

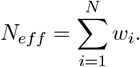

The measure depends on the choice of similarity metric and threshold, which are typically heuristic. *N*_*eff*_ is widely used to mitigate sampling bias in MSAs, particularly in profile construction and coevolutionary analyses. However, it reflects only the number of non-redundant sequences under a chosen similarity criterion, rather than the diversity implied by the underlying sequence distribution from which the MSA is sampled.

### Average pairwise distance

Average pairwise distance summarizes diversity through a fraction of mutations *d*_*ij*_ between sequences *i* and *j*. It is defined as

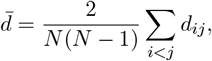

where *N* is the number of sequences. This quantity has a direct statistical interpretation as the expected distance between two sequences drawn uniformly at random from the alignment. Therefore, it implicitly assumes that the distribution of pairwise distances is approximately Gaussian or unimodal. Average distance is often used as a coarse measure of global divergence. Its main limitation is that it becomes uninformative when clustered or multimodal structure is present, because averaging small intra-cluster distances with large inter-cluster distances can yield a mean that does not correspond to any actual sequence relationship.

## Results

### Definition of *L*_*eff*_

Now I describe *L*_*eff*_ . Sequences of length *L* over an alphabet of size *k* are encoded as indicator vectors called one-hot (see Methods), giving *X* ∈ *R*^*N*×*kL*^. Because each *k* symbol block satisfies a sum-to-one constraint, it contains only (*k* − 1) independent degrees of freedom. To work directly in a non-redundant space, each block is projected onto a (*k* − 1) dimensional zero-sum basis *Q* using Helmert’s contrastive encoding [12].

Each sequence is given a weight 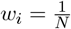 where ∑_*i*_ *w*_*i*_ = 1 which is embedded in *W* = diag(*w*_1_, …, *w*_*N*_). Weighted centered features are 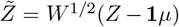, with weighted mean *µ* = **1**^⊤^*WZ*, and the covariance matrix is 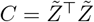.

The eigenvalues of *C, λ*_*j*_, are normalized as *p*_*j*_ = *λ*_*j*_*/* ∑_*ℓ*_ *λ*_*ℓ*_. Because *C* is symmetric and positive semidefinite, the eigenvalues are nonnegative. The spectral entropy

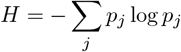

quantifies how broadly the observed sequence variation is distributed across the alignment. In an MSA, a high value indicates that many different patterns of variation are present across the sequences, whereas a low value indicates that the variation is concentrated in only a few recurring patterns.

I convert this spectral entropy into an effective length,

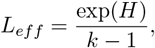

which is the effective dimensionality (or effective rank) of the MSA matrix divided by the alphabet size [13]. This quantity is interpreted as the length of a hypothetical alignment in which every position varies independently and with the full alphabet, yet produces the same total amount of variation as the observed MSA. From this, an effective support size is obtained as

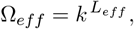

representing the number of distinct sequence configurations effectively supported by the data.

The computation of *L*_*eff*_ is highly efficient as it scales with the feature dimension *k* × *L* rather than the sample size *N*, allowing for the processing of typical MSAs (*L* = 200) in a fraction of a second. In the data-poor regime (*N* ≪ *kL*), the framework preserves this efficiency by exploiting the duality of the eigenvalue spectrum: the covariance between sequences yields the identical set of non-zero eigenvalues as the feature covariance matrix because they share the same singular values. The approach is implemented in Python code available at https://github.com/vaiteaopuu/effective_length. For MSAs with large *kL*, I leverage the Singular Value Decomposition (SVD) of 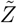 so that it does not require the explicit calculation of the covariance matrix, which drastically reduces computation time, see Fig SI1.

To quantify the overlap between the sequence spaces spanned by two independent MSAs *X* and *Y*, I define the cross effective length 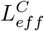. After weighted centering of both alignments, the cross operator is

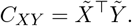

The singular value spectrum of *C*_*XY*_ is processed through the same spectral–entropy pipeline used for *L*_*eff*_, yielding an effective dimensionality that measures the alignment of their principal modes of variation. A large 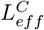 indicates that the two datasets explore similar regions of sequence space, whereas small values reflect weakly overlapping variability. For comparability across datasets, I report the normalized quantity

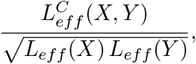

which ranges from low overlap (near 0) to near-complete alignment (approximately 1).

In conclusion, *L*_*eff*_ is a classic measure of information that is expressed in sequence length for interpretability. This framework is formally analogous to Principal Component Analysis (PCA) or SVD [14]. However, the method distinguishes itself technically by employing zero-sum (Helmert) encoding to resolve the one-hot limitations [15, 2, 14]. I show below that this encoding is critical to removing the one-hot bias. Moreover, this approach is not equivalent to simply counting how many modes explain an arbitrary variance threshold; rather, spectral entropy acts as a continuous, parameter-free summary of the effective dimensionality by weighting the contribution of the entire eigenvalue spectrum, including the tail.

### Application of *L*_*eff*_ to synthetic MSAs

I now examine how *L*_*eff*_ responds to changes in global variation under controlled conditions. To do so, I generated synthetic MSAs from a multivariate Gaussian model defined in one-hot space, where the strength of correlation across positions can be tuned directly through a single parameter *ρ* (see Methods).

I first generated MSAs of length *L* = 10 over an alphabet of size *k* = 4, using sample size *N* = 100 and varying *ρ* from 0.1 to 1.0. At *ρ* = 1.0, all positions are strongly correlated in a way that only sequences with the same symbol are sampled. For each value of *ρ*, I produced 30 independent MSAs and computed four diversity measures: the effective length *L*_*eff*_, positional entropy, 3-mer entropy, the average distance, and the effective number of sequences *N*_*eff*_ . To facilitate the representation in the result figure, I converted positional entropy into perplexity which is the average positional entropy exponentiated. The results are shown in Fig 2. As *ρ* increases, the model produces increasingly constrained alignments. This collapse of global variation was captured clearly by *L*_*eff*_, 3-mer entropy, and *N*_*eff*_, all of which decreased with increasing correlation. Their rates of decrease differed, reflecting their distinct sensitivities: *L*_*eff*_ showed a smooth monotonic decline that tracked the reduction in global variability; k-mer entropy decreased more gradually because it captures only local motif diversity; and *N*_*eff*_ decreased only once pairwise sequence identities crossed the similarity threshold (1-0.8=0.2) used to define redundancy. In contrast, positional entropy and average distance remained nearly constant for all *ρ*. Positional entropy remains constant because per-site symbol frequencies do not change under the chosen covariance matrix. For the average distance, the convergence into increasingly similar clusters is compensated by the higher intercluster distance (see Fig SI4).

**Figure 1:**
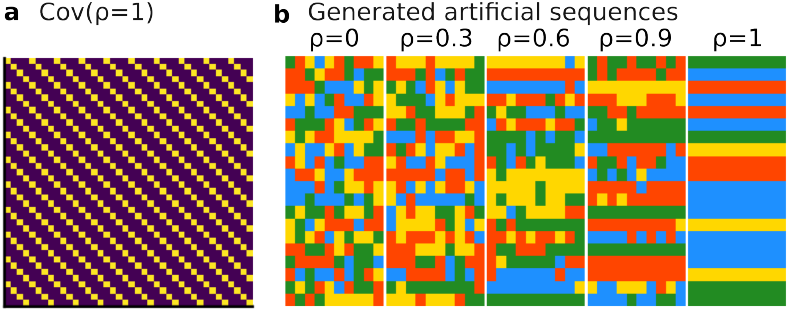
Synthetic MSA generation. **(a)** Visualization of the covariance matrix for the generative model with the coupling parameter set to *ρ* = 1 (fully coupled). Yellow entries indicate a correlation of 1, while other entries are 0. **(b)** Representative synthetic sequences generated with varying coupling strengths *ρ*, ranging from independent sites (*ρ* = 0) to fully coordinated motifs (*ρ* = 1). This illustrates the transition from random noise to structured patterns.

**Figure 2:**
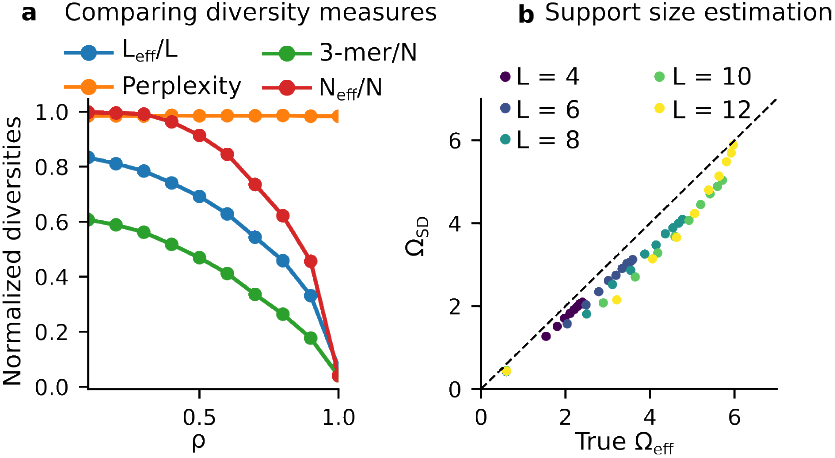
Benchmarking diversity measures on synthetic data. **(a)** Normalized diversity metrics (*L*_*eff*_ */L*, 3-mer entropy, Perplexity/*k*, and *N*_*eff*_ */N*) plotted as a function of the coupling parameter *ρ. L*_*eff*_ tracks global variation, which decays smoothly. **(b)** Scatter plot of the estimated effective support size 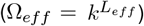 versus the true support size of the generative multivariate Gaussian model, colored by sequence length *L* (4 to 12). The dashed line represents the identity function (*y* = *x*).

I next assessed whether *L*_*eff*_ can recover the support size of the underlying multivariate Gaussian generative model. For each sequence length *L* ∈ {4, 6, 8, 10} and correlation value *ρ*, I approximated the true support by sampling a large number of sequences (10^5^) and computing an empirical entropy of the resulting distribution. From much smaller samples of size *N* = 100, I estimated 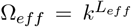. Across all lengths and correlations, Ω_*eff*_ closely matched the ground-truth support size, lying near the identity line on a log scale, see Fig 2. I performed the same analysis using only the one-hot encoding, which resulted in a biased estimation, see Fig SI6. The latter demonstrates that the Helmert’s encoding is critical to achieve these results. Independent-site estimates derived from positional entropy consistently overestimated support in correlated conditions, whereas the estimate derived from *L*_*eff*_ remained accurate even when correlations were strong. *k*-mer entropy is not applicable in this setting because its estimate cannot be mapped to full-length sequence support. Similarly, *N*_*eff*_ does not provide a support estimate, as it reflects redundancy reduction rather than the size of the underlying sequence distribution.

Together, these results show that *L*_*eff*_ provides a consistent measure of global variation across sequence space, thereby offering a complementary measure of diversity. Applied to real MSAs, it should provide an accurate estimate of the diversity implied by the observed covariance structure.

### Diversity of RNA MSAs

#### Diversity and predicted effective size of RNAs

I now measure the diversity of real RNA MSAs, which I chose because independent support-size estimates are also available (obtained with direct coupling analysis (DCA) [15]). These MSAs are described in more detail in the Methods.

Across the 25 curated RNA MSAs, the effective length (*L*_*eff*_) was consistently much smaller than the alignment length (L), with an average *L*_*eff*_ of 28.18 (see Fig 5), corresponding to a 4.5-fold reduction relative to the mean MSA length (*L*_*eff*_ /*L* = 0.23). This indicates strong evolutionary and structural constraints. Notably, this value is substantially lower than what is expected from secondary-structure constraints alone. For instance, generating 500 sequences constrained to the tRNA fold using RNAinverse [16] yielded *L*_*eff*_ = 37.37, whereas the natural tRNA MSA (∼ 30, 000 sequences) extracted from [15] produced *L*_*eff*_ = 21.2. RNAinverse is a method that generates sequences predicted to adopt a specified target secondary structure. Extending this comparison to all 25 families, sequences generated solely from their consensus secondary structures (computed with ViennaRNA [16]) yielded an average *L*_*eff*_ = 54.58, still 2.23-fold below the alignment length but approximately twice the diversity observed in natural MSAs.

In contrast, generative models populate markedly different volumes of sequence space. Libraries sampled from DCA models for the same families exhibited a much higher average *L*_*eff*_ = 64.93, suggesting that such models can explore a sequence space approximately 2.48 times larger than natural diversity, see Fig 5. By comparison, sequences generated with a variational autoencoder (VAE) yielded *L*_*eff*_ = 27.26, closely matching the natural MSAs. These results indicate that models trained on the same data can produce substantially different effective sequence spaces. However, higher diversity does not necessarily imply functionality; this aspect is examined in the following section on experimental ribozyme libraries.

I further assessed whether *L*_*eff*_ provides a meaningful estimate of support size by comparing 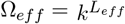 with independent support-size estimates obtained from a DCA variant (eaDCA) [15] across the same 25 RNA families. Using full MSAs, the log–log relationship between the two measures yielded a Pearson correlation of *ρ* = 0.79 (Fig 5). Applying the classical redundancy reweighting implemented in [10] increased the correlation to *ρ* = 0.92, and adding regularization (*α* = 0.01) further improved it to *ρ* = 0.94.

Finally, I observed a connection between *L*_*eff*_ and the RNA folding thermodynamics. This diversity measure corresponds to the length of a fully random MSA, which can be understood as the alignment of an RNA that is completely unfolded. In such a state, the molecule contributes only configurational entropy, which is proportional to its length. I tested this interpretation using the MSAs and comparing *L*_*eff*_ with the folding free energy of the consensus secondary structure predicted by `RNAalifold` [16], obtaining a Spearman correlation of *ρ* = 0.6 (p-value = 0.003, shown in Fig 4). Using sequences generated using RNAinverse on randomly generated folds of exactly the same length but varying stability (controlled by the number of paired nucleobases), I obtained an even higher correlation of *ρ* = 0.7 (see Fig SI5). These results provide evidence of the proposed connection.

Taken together, these results show that natural RNA families explore only a fraction of the sequence space compatible with their secondary structure, while generative models can access broader regions of this space. The observed association with folding stability further establishes the interpretability of *L*_*eff*_ .

#### Diversity of experimental library of RNA sequences

I now measure the diversity of experimentally generated RNA sequence libraries by analyzing the comprehensive dataset produced in [2]. In this study, multiple generative modeling approaches were used to design large sets of the group I intron of the Azoarcus bacterium, which were subsequently synthesized and assayed for catalytic activity. Here, I analyze the diversity of the active sequences only unless otherwise mentioned (5895 sequences). The resulting collection provides a unique opportunity to evaluate how different computational design pipelines populate sequence space and how their diversity changes as variants accumulate mutations away from the wild type. As high-throughput experimental screens of model-generated sequences are becoming increasingly common, establishing a principled way to quantify and compare the diversity of such libraries is essential. Here, I omit *k*-mer entropy, whose interpretation is unclear, and N_eff_, which in this regime either remains close to one when normalized or simply reflects the number of sequences in each bin. All results are shown in Fig 3.

**Figure 3:**
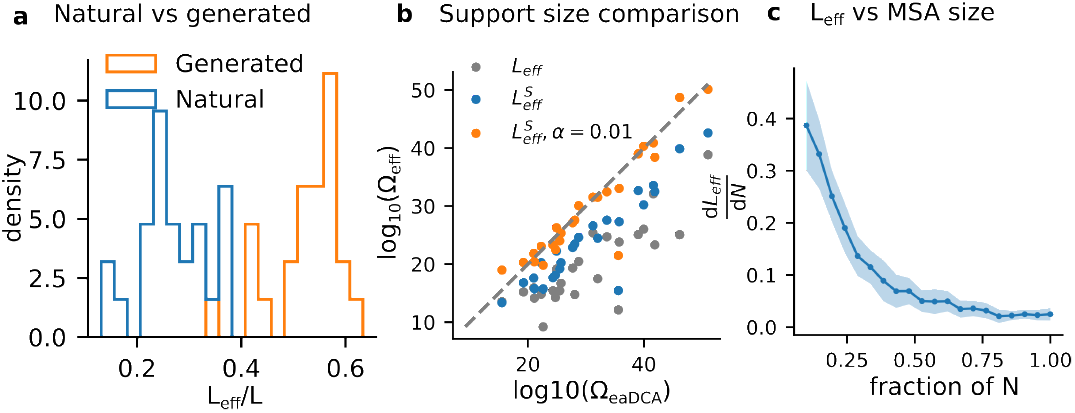
Comparison of Natural vs. generated MSAs and support size validation. **(a)** Density histogram showing the distribution of the normalized effective length (*L*_*eff*_ */L*) for the 25 natural RNA MSAs (blue) and DCA generated libraries (orange). **(b)** Scatter plot of the spectral support size estimate (Ω_*eff*_) versus the independent estimate (Ω_*eaDCA*_) on a log_10_ scale. Grey dots indicate support sizes calculated using the full MSA. Blue dots indicate *L*_*eff*_ support sizes calculated by reweighting sequences with the the redundancy reduction derived from *N*_*eff*_ [10]. Orange dots indicate the *L*_*eff*_ with redundancy removed and regularization. **(c)** Diversity saturates as the size of MSA increases. For each RNA MSA, I subsample a fraction of the sequences and computed *L*_*eff*_ . I then evaluated the marginal gain in diversity as a function of sample size, quantified by 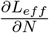. The curve shows the average across all RNA families (solid line), and the shaded region indicates the 95% confidence interval.

**Figure 4:**
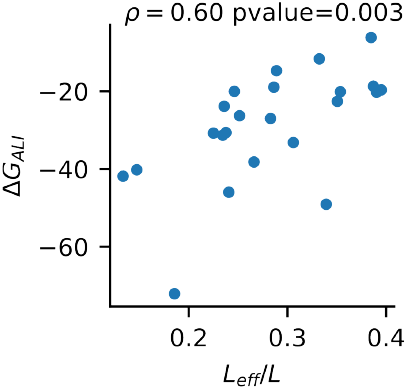
Physical interpretation of effective length. Scatter plot showing the relationship between the consensus folding free energy (Δ*G*_*ALI*_ in kcal/mol) on the y-axis and the normalized effective length (*L*_*eff*_ */L*) on the x-axis for 25 RNA families. The plot reports the Spearman rank correlation (*ρ* = 0.60) and the associated p-value (0.003).

**Figure 5:**
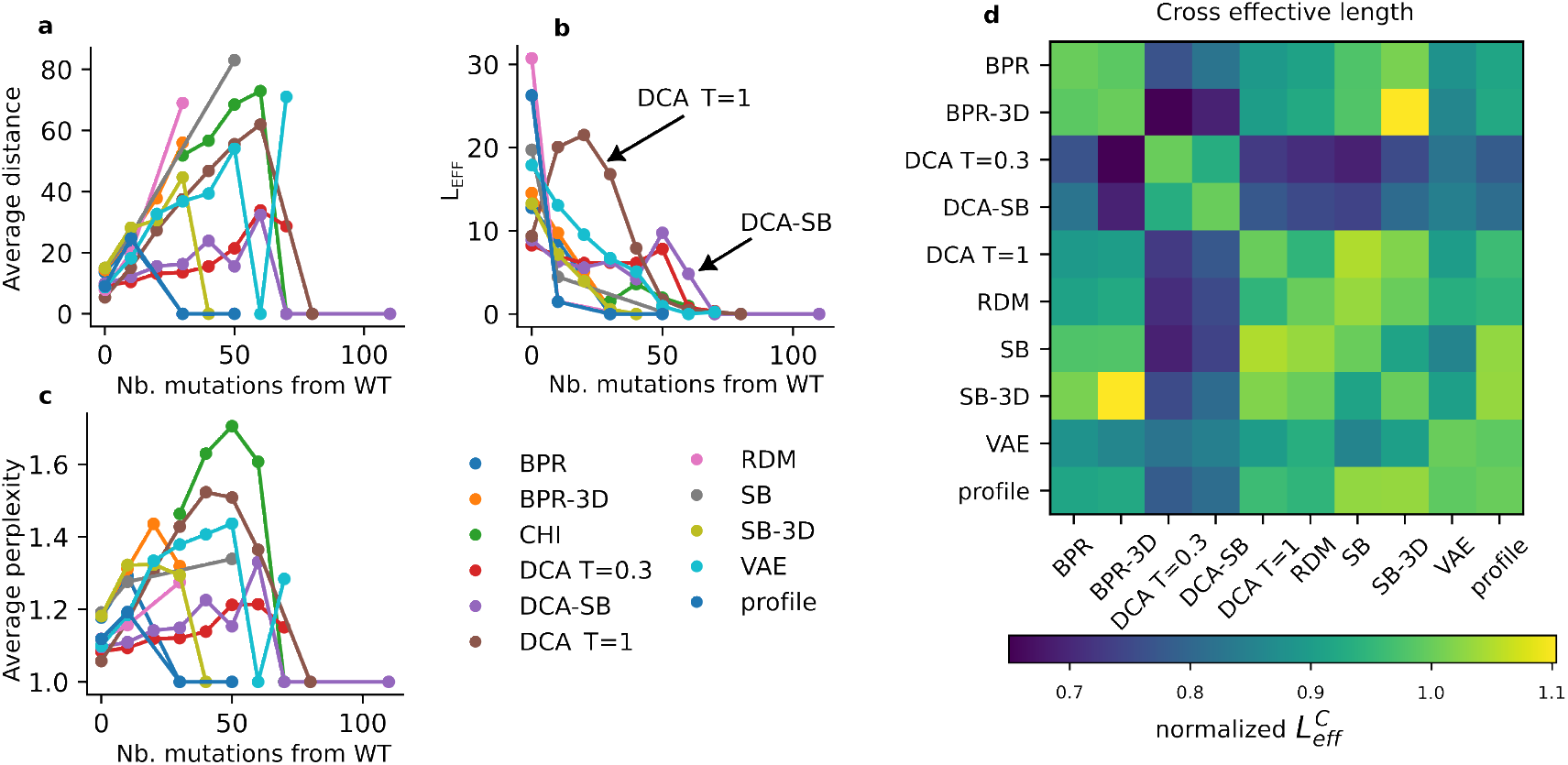
Diversity of experimentally generated RNA libraries. Diversity metrics plotted against the number of mutations from the wild type (WT) sequence for various generative models. **(a)** Average pairwise Hamming distance between sequences. **(b)** The effective sequence length (*L*_*eff*_). **(c)** Average positional perplexity. The different colored lines correspond to the specific generative models and baselines (e.g., DCA, VAE, RDM, CHI) listed in the legend [2]. **(d)** Estimated overlap between generated libraries. Heatmap of the normalized cross effective length 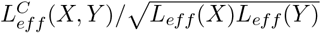 computed between RNA sequence libraries obtained with different generative processes. Warmer colors denote stronger overlap. To allow the comparison between all models, I restricted the analysis to the functional sequences with 20 mutations from the wild type.

#### Average pairwise distance

The distance grows approximately linearly with the number of mutations from the wild type, indicating that as mutations accumulate, the generated functional sequences become increasingly dissimilar. RDM reaches the largest values among methods, as it samples mutations fully at random (uniformly across positions and nucleotides). However, this result is unexpected because only a very small number of those variants were functional. The average pairwise distance is sensitive to clustered structure and implicitly assumes a unimodal distance distribution: a small number of well-separated functional clusters can yield a large average even if within-cluster diversity is low. Moreover, removing sequences that interpolate between clusters can increase the mean, while adding a distant sequence that forms a new cluster may leave it nearly unchanged, even though the underlying structural diversity has increased. Consequently, this measure provides a misleading picture of diversity because the mean distance is not monotonic with the true geometric organization of the dataset.

#### Positional entropy

To make this measure more interpretable, I converted positional entropy to perplexity, given by the exponential of the average site-wise entropy. This quantity reaches its maximum at approximately 50 mutations in the CHI dataset, which is derived from natural homologs with indels replaced by the wild-type nucleotide. This result is inconsistent with the average pairwise distance. In structured RNAs, however, sequence variation is strongly constrained by conserved stems, loops, and long-range base-pairing interactions. Because positional entropy treats sites independently, it necessarily ignores these structural constraints and therefore counts variability at positions that cannot vary freely in combination.

#### Effective sequence length *L*_*eff*_

*L*_*eff*_ captures global diversity and therefore separates model behaviours more clearly. Most approaches exhibit a steady decrease in *L*_*eff*_ as mutation counts rise, which is consistent with the fact that only a narrow subset of highly mutated variants remains functional as mutations accumulate. DCA achieves the highest *L*_*eff*_ over intermediate mutation ranges. At larger mutational distances (≥50), the hybrid DCA–SB model maintains higher values. The VAE, despite recovering functional sequences at rates comparable to DCA, produces noticeably lower diversity. This result is consistent with the reduced diversity observed in the above section for VAE.

Generative models can differ not only in the volume of the functional sequence space they explore, but also in the specific regions they occupy within the RNA landscape. Figure∼3 reports the normalized cross effective length between all pairs of libraries, enabling a direct comparison of their relative overlap. For this analysis, I restricted the sets to functional sequences below 20 mutations from the wild type. DCA *T* = 0.3 and DCA–SB cluster closely, consistent with their shared Potts parameters and identical MCMC sampling procedure, whereas DCA *T* = 1, derived from the sparse eaDCA variant, spans a more distinct region. BPR and BPR-3D are similarly aligned, differing only by fixing tertiary-interaction positions to the wild-type nucleotide; the same correspondence is recovered between BPR-3D and SB-3D, which constrain identical sites. These relationships are not detected by cross average-distance analyses (Fig SI7), highlighting the ability of the cross effective length to capture global geometric similarity between generative designs and existing data.

Generative models with similar functional yields can occupy very different volumes of functional sequence space. Average pairwise distance systematically overestimates diversity because a few distant clusters inflate the mean despite low within-cluster variability. Positional entropy is more stable but ignores long-range interactions, which inflates diversity when coordinated constraints are present. *L*_*eff*_, by incorporating global correlations, provides a more faithful estimate of the effective functional diversity explored by each design strategy. Moreover, 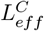 allows one to compare two sets of related sequences to determine whether they overlap.

### Diversity of protein MSAs for fold prediction

I next test if the MSA diversity is related to the protein structure prediction performance with four state-of-the-art deep learning models: AlphaFold [4], RoseTTAFold [17], ESMFold [18], and OmegaFold [19]. This analysis utilized protein MSAs from the study [20], focusing on the correlation between the diversity *L*_*eff*_ and two key quality metrics: the Local Distance Difference Test (LDDT) [4] and the Template Modeling Score (TM-score) [21].

Across the 60 analyzed protein families, the average is *L*_*eff*_ = 17.1, which is lower than the diversity observed in RNA. The latter cannot be explained solely by the larger amino acid alphabet (see Fig SI3). The average ratio is *L*_*eff*_ /*L* = 0.07. To confirm this result, I computed *L*_*eff*_ across 1000 MSAs taken randomly from the Uniclust30 database [22] which contains one MSA per cluster of 30% identity defined in the UniProt database. The obtained average *L*_*eff*_ = 21.8. This result confirms that the evolutionary constraints applied on proteins are typically stronger than for RNAs.

While the sequences used to train structure prediction models are essentially the same, the ways in which they are used vary drastically: in AlphaFold and RoseTTAFold, the model input is an MSA; in ESMFold and OmegaFold, the input is a single sequence. Despite these differences, a positive correlation is observed between *L*_*eff*_ and model TM-scores across all platforms except AlphaFold. In contrast, no significant correlation was found between the number of sequences *N* per MSA (or sequence length *L*) and prediction accuracy. Similarly, *N*_*eff*_ did not show any significant correlation either, which confirms the results published earlier in [11]. For LDDT, correlations were stronger for all models except AlphaFold. *N*_*eff*_ displayed significant but weak associations. These results suggest that *L*_*eff*_ could help in anticipating how challenging a prediction is.

The association strength with MSA diversity varies across the two types of models. For LDDT and TM-score, single-sequence models exhibited the strongest associations, with Spearman correlations of *ρ* = 0.58 for ESMFold and *ρ* = 0.54 for OmegaFold. I chose Spearman because there is no reason to believe that the relation between performance and diversity is linear. RoseTTAFold followed with *ρ* = 0.50, while AlphaFold showed a more moderate correlation of *ρ* = 0.33. The global fold accuracy, measured by TM-score, followed a similar trend; ESMFold and OmegaFold demonstrated the most robust connection to sequence diversity (*ρ* = 0.452 and *ρ* = 0.539 respectively), whereas AlphaFold showed no significant correlation (p-value = 0.233). These benchmarks were performed on the CASP15 dataset [20]. Although the amount of information available is similar, their performances from single-sequence-based to MSA-based models vary notably.

To assess whether a protein family contains sufficient evolutionary information to support accurate structure prediction, I estimated model-specific *L*_*eff*_ thresholds based on LDDT. I define high local accuracy as LDDT ¿ 0.7, consistent with commonly used confidence interpretations in [4]. For each predictor, I determined a cutoff *τ* such that families with *L*_*eff*_ ≤ *τ* satisfy P(LDDT > 0.7) < 0.5. Although some architectures operate on single sequences, an auxiliary MSA was still constructed solely to quantify family diversity; the alignment itself was not used as model input. This procedure provides a criterion to estimate *a priori* whether a case is expected to be accurately predicted. The resulting thresholds were *τ* ≈ 18.69 for ESMFold, *τ* ≈ 4.96 for AlphaFold, *τ* ≈ 26.78 for OmegaFold, and *τ* ≈ 37.27 for RoseTTAFold.

Prediction accuracy is well captured by *L*_*eff*_ . The strong correlation between *L*_*eff*_ and the performance of single-sequence models, together with the weaker dependence observed for MSA-based models, suggests that alignment procedures further enhance the evolutionary signal used for protein structure prediction. These results provide an operational threshold of diversity to estimate the accuracy in structure predictions.

## Discussion

This work introduces an information-theoretic framework to quantify sequence diversity in multiple sequence alignments through the effective sequence length, *L*_*eff*_ . In addition to its interpretability in sequence length, it differs from PCA-based measures used so far because it relies on a zero-sum projection that is critical for the accuracy of it. By leveraging spectral entropy, *L*_*eff*_ provides a measure of global variation that captures correlations distributed across positions, rather than local or pairwise summaries. Synthetic benchmarks demonstrated that *L*_*eff*_ reliably tracks reductions in accessible sequence space induced by increasing constraints, whereas commonly used measures such as positional entropy or average pairwise distance fail to do so under correlated constraints.

Applied to natural RNA families, *L*_*eff*_ reveals that RNA MSAs occupy a highly restricted region of sequence space, with an average effective length of 28.18 positions, approximately 4.5-fold smaller than the alignment length. This constraint is substantially stronger than what is expected from secondary structure alone, as shown by comparisons with sequences generated to satisfy identical folds. This indicates that evolutionary selection imposes strong constraints beyond base pairing, likely reflecting requirements on folding kinetics, tertiary contacts, and functional robustness. The observed correlation between *L*_*eff*_ and folding free energy further supports a physical interpretation of *L*_*eff*_ as a proxy for configurational entropy, linking sequence diversity to thermodynamic stability.

Proteins exhibit even smaller effective lengths than RNAs, despite their larger alphabet, indicating even stronger evolutionary and structural constraints. Across protein families, structure prediction accuracy correlates with *L*_*eff*_ but not with the raw number of sequences in the MSA. This result shows that model performance scales with informative diversity rather than dataset size or effective sequence count. The particularly strong dependence observed for single-sequence models suggests that MSA-based methods implicitly amplify evolutionary information, partially compensating for limited intrinsic diversity. In contrast, when such amplification is absent, prediction accuracy becomes directly constrained by the effective richness of the underlying sequence family. Building on these observations, I propose empirical diversity thresholds to estimate the minimum *L*_*eff*_ required to reliably achieve high structural prediction accuracy. These thresholds can readily be used to anticipate the prediction accuracy to expect.

The analysis of experimentally generated ribozyme libraries illustrates the utility of *L*_*eff*_ for evaluating generative models, particularly in light of the limitations of commonly used measures such as average pairwise distance and positional entropy, which respectively overestimate diversity through cluster separation and ignore long-range constraints. To complement *L*_*eff*_, I also introduced a cross effective length 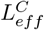 measure that quantifies the overlap between the sequence spaces explored by two different samples, providing, to my knowledge, the first model-based measure of inter-library similarity. This analysis reveals that covariance-based models explore a broader functional sequence space than variational autoencoders, even when both achieve comparable fractions of active sequences. Consequently, *L*_*eff*_ can serve as an operational tool to guide sequence design toward underexplored regions of sequence space, for example within active learning frameworks.

More broadly, expressing diversity on an effective-length scale enables quantitative decision rules for sequence datasets. *L*_*eff*_ can be used to set minimum diversity thresholds before training predictive models, to stop data collection once additional sequences no longer increase effective information, and to select new designs that maximize diversity gain rather than raw count. In iterative or active-learning pipelines, it can serve directly as an objective function to bias sampling toward underrepresented regions of sequence space. Because it is model-agnostic and comparable across datasets, the same criterion can guide curation, experimental allocation, and cross-library comparison whenever correlations dominate variability.

## Methods

### One-hot

Sequences are represented using a standard one-hot encoding scheme to map discrete symbols into a vector space. A sequence of length *L* over an alphabet of size *k* (e.g., *k* = 5 for RNA, including the gap) is encoded as a binary vector **x** ∈ {0, 1} ^*kL*^. For each position *i* ∈ {1, …, *L*}, the state is represented by a block of *k* binary variables, where the entry corresponding to the observed symbol is set to 1 and all others to 0. An MSA containing *N* sequences is therefore represented as a binary matrix *X* ∈ {0, 1} ^*N*×*kL*^, where each row corresponds to a single encoded sequence.

### Helmert encoding of MSAs

Helmert encoding replaces categorical one-hot vectors with an orthonormal contrast basis that removes the linear dependence inherent in one-hot representations [12]. As an example, I consider here RNA sequences with four-nucleotide alphabet {*A, C, G, U*}. The encoding uses the orthonormal contrast matrix:

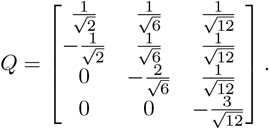

The columns of *Q* span the zero-sum subspace and are orthonormal, yielding successive contrasts: first comparing *A* with *C*, then the mean of {*A, C*} with *G*, and finally the mean of {*A, C, G*} with *U* . Multiplying each one-hot nucleotide vector by *Q* maps it to a three-dimensional orthonormal coordinate, preserving all information except the redundant overall offset. Applying this transformation along an RNA sequence produces a compact and statistically well-conditioned representation suitable for downstream modeling. This procedure is generalizable to amino acids as well. For MSAs, I include gaps as an additional symbol.

### Synthetic MSA generation

To produce synthetic MSAs with controlled amounts of coordinated variation, we construct a covariance directly in one-hot space and tune a coupling parameter *ρ*. At each position, I begin with the standard categorical covariance, which captures how symbols fluctuate relative to one another at a single site. To introduce dependence across positions, I mix this position-wise structure with a fully coupled component. In this coupled term, a given symbol (e.g. “A”) at one position co-varies only with the same symbol at all other positions and is anticorrelated with the other symbols. As *ρ* increases, the model increasingly favors configurations in which many positions simultaneously shift toward the same symbol. When *ρ* is large, the resulting alignments are dominated by sequences that are composed mostly of a single symbol pattern, whereas small *ρ* yields nearly independent site variation. Gaussian samples drawn from this covariance are decoded blockwise by selecting the largest component, producing categorical sequences that reflect the intended level of global coordination. The specific algorithm to sample sequences from the covariance matrix is described in supplementary information.

### Natural RNA MSAs

To estimate available diversity in natural MSAs, I extracted RNA families from one study. I incorporated the 25 RNA families from [15] for which independent estimates of effective support size were previously obtained. The average MSA size is 3799 sequences. This dataset enables a direct comparison between *L*_*eff*_ and established model-based estimates of variability. Together, these two sources provide a broad and structurally grounded benchmark for quantifying how much meaningful variation natural RNA families contain.

I constructed an experimental benchmark using data from a high-throughput study in which multiple generative modeling approaches were used to design variants of the Azoarcus ribozyme (197 nucleotides) and experimentally measure their activity. The study evaluated a broad spectrum of sequence-generation methods, including simple baselines—random uniform mutagenesis (RUM) and independent profile sampling (PRO)—structure-aware mutational schemes (BPR, BPR-3D, SB, SB-3D), evolution-based statistical models derived from natural intron alignments (DCA sampled at *T* = 1 and *T* = 0.3), a variational autoen-coder trained on the same data (VAE), a hybrid Potts–structure approach (DCA-SB), and a reference set of chimeric natural introns (CHI). For each of these design strategies, I extracted all sequences that were synthesized and experimentally assayed together with their measured activities at 37^°^C. After applying the quality and filtering criteria used in the original screen, this yielded a comprehensive dataset of designed ribozyme variants paired with quantitative activity measurements, which I use to evaluate how diversity measures relate to only functional sequences. The dataset is composed of 15146 entries where 5895 were found active (0.39%).

## Acknowledgment

I thank Philippe Nghe and Martin Weigt for useful discussions.

## Code and data availability

The code to reproduce all analyses is available at https://github.com/vaiteaopuu/effective_length. RNA MSAs and support-size estimates were extracted from [15]. Ribozyme sequences and catalytic activities were taken from [2]. Protein MSAs were randomly sampled (n = 1000) from [22]. MSAs used for structure-prediction benchmarks were obtained from [20].

**Table 1:** Effect of sequence diversity in structure prediction. Spearman correlation between *N*_*eff*_, *L*_*eff*_, *N*, and *L* with TM-score and LDDT (associated p-values) across structure prediction models.

| Method | $L_{eff}$ | $N_{eff}$ | $N$ | $L$ |
| --- | --- | --- | --- | --- |
| OmegaFold | TM 0.539 (5.23e-6) | TM 0.243 (0.0549) | TM 0.0995 (0.438) | TM -0.196 (0.124) |
|  | LDDT 0.676 (1.20e-9) | LDDT 0.449 (2.23e-4) | LDDT 0.253 (0.0457) | LDDT -0.112 (0.382) |
| ESMFold | TM 0.452 (2.03e-4) | TM 0.191 (0.134) | TM 0.0696 (0.588) | TM -0.219 (0.0849) |
|  | LDDT 0.619 (6.51e-8) | LDDT 0.397 (0.00126) | LDDT 0.178 (0.164) | LDDT -0.206 (0.106) |
| RoseTTAFold | TM 0.305 (0.0151) | TM 0.0273 (0.832) | TM -0.0458 (0.721) | TM -0.184 (0.150) |
|  | LDDT 0.592 (3.25e-7) | LDDT 0.367 (0.00312) | LDDT 0.198 (0.120) | LDDT -0.130 (0.311) |
| AlphaFold | TM 0.152 (0.233) | TM -0.0339 (0.792) | TM -0.0851 (0.507) | TM -0.109 (0.397) |
|  | LDDT 0.361 (0.00369) | LDDT 0.129 (0.315) | LDDT 0.0441 (0.732) | LDDT -0.118 (0.357) |

## SI

### Sampling sequences from a multivariate Gaussian model

How I sampled sequences from a covariance matrix *C. C* is the covariance matrix of a one-hot encoded MSA. To sample sequences from this distribution assuming an average symbol distribution *µ* = (1*/k*, …1/*k*). First, we compute the eigenvalues and eigenvectors (*λ*_*i*_, *v*_*i*_) of *C*. The second step is to sample a novel sequence 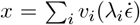), where *ϵ* ∼ 𝒩 (0, 1). The last step consists in converting the *x* vector into a one hot encoded sequence with nearest-neighbor approach which assigns the symbol identity of the closest one hot encoded nucleotide. Since we assume a uniform distribution of the symbols, the positional perplexity is always maximized, regardless of the interaction level encoded in the covariance matrix.

### Direct Coupling Analysis

Direct Coupling Analysis (DCA) infers residue–residue interactions by fitting a pairwise maximum-entropy (Potts) model that reproduces the empirical single-site and pairwise frequencies observed in a multiple sequence alignment [10]. The statistical model is

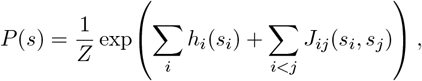

where *s* = (*s*_1_, …, *s*_*L*_) denotes a sequence, *h*_*i*_ are site-specific fields, *J*_*ij*_ are direct couplings, and *Z* is the partition function. The inferred couplings quantify conditional dependencies that remain after removing indirect correlations, enabling high-resolution contact prediction and structural inference. Extensions such as eaDCA have been applied to estimate the effective support size of high-dimensional sequence distributions [15], and DCA-based generative models have been used for rational sequence design, including the *in silico* design of functional RNAs [2]. To run DCA (fully connected), I adapted the python code from [23] which runs Gibbs Markov Chain Monte Carlo for 100 sweeps. We checked the convergence by comparing pairwise frequencies between a 1000 MCMC sample and the MSA.

### Variational autoencoder

The variational autoencoder (VAE) provides a generative model for RNA sequence families by learning a latent representation from multiple sequence alignments [24]. Each sequence is mapped to a continuous latent variable drawn from a Gaussian distribution, and a neural decoder transforms this latent vector into position-wise categorical probabilities over the nucleotide alphabet. Training maximizes the evidence lower bound, combining a reconstruction term with a KL regularizer that encourages the inferred latent distribution to remain close to a standard normal prior [25]. To train the models, I ran 100 epochs (with batch size 32) of learning using the Adam optimizer [26]. After training, new sequences are generated by sampling latent vectors from the prior, *z* ∼ 𝒩 (0, *σ*^2^*I*), and decoding them into categorical logits at each sequence position; discrete sequences are then produced by selecting the most likely nucleotide at each site. For our experiments, we typically generate 1000 sequences.

**Figure SI1:**
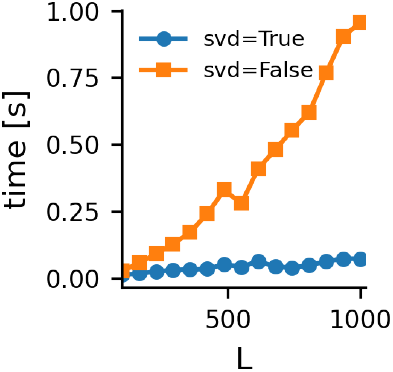
Runtime of the effective length computation as a function of sequence length *L* for fixed sample size *N* = 100 and alphabet size *k* = 4. Synthetic MSAs are generated from a Gaussian covariance model with correlation parameter *ρ* = 0.5. Execution time is reported for two implementations: singular value decomposition and covariance eigen-decomposition. Each point corresponds to the mean wall–clock time over independent realizations.

**Figure SI2:**
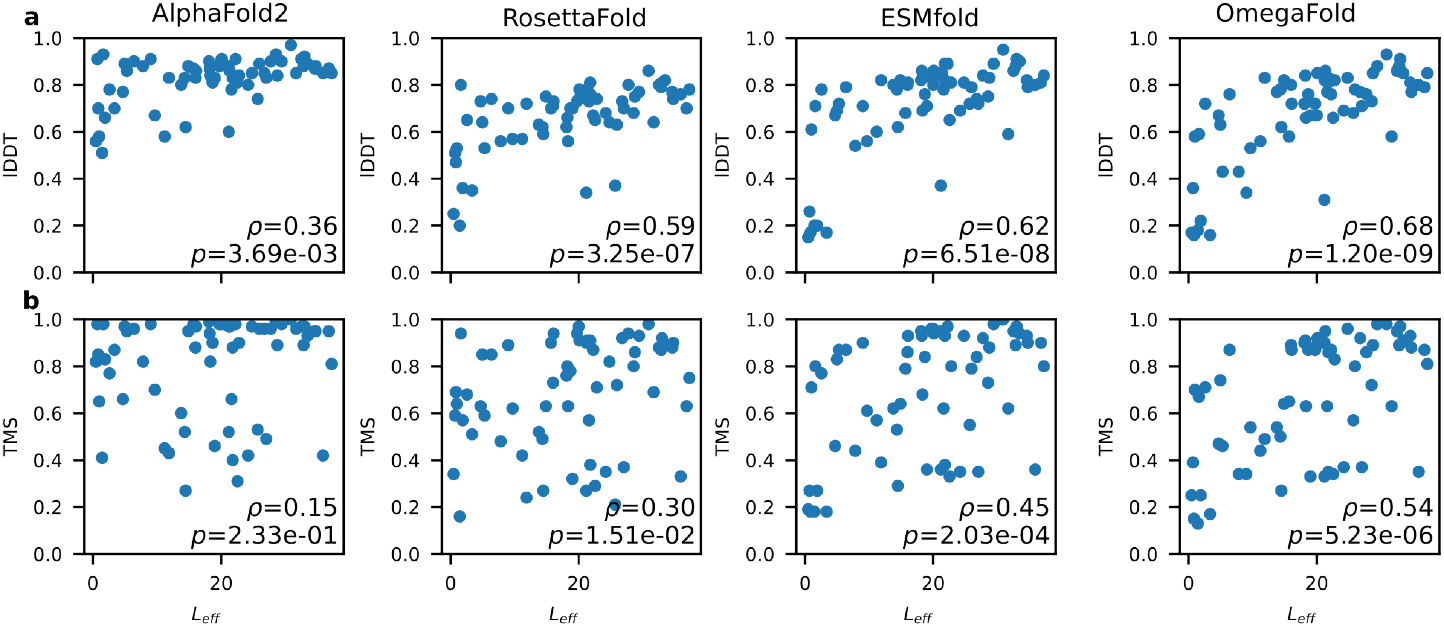
Relationship between sequence diversity (*L*_*eff*_) and protein structure prediction quality. **(a)** Scatter plots showing the correlation between the effective length (*L*_*eff*_) and the Local Distance Difference Test (LDDT) for four transformer-based models: AlphaFold2, RoseTTAFold, ESMFold, and OmegaFold. Spearman correlation coefficients (*ρ*) and p-values are shown for each model, with single-sequence PLM-based models (ESMFold and OmegaFold) exhibiting the strongest dependencies on family diversity. **(b)** Corresponding correlations for the global TM-score. All benchmarks were performed on the CASP15 dataset [20].

**Figure SI3:**
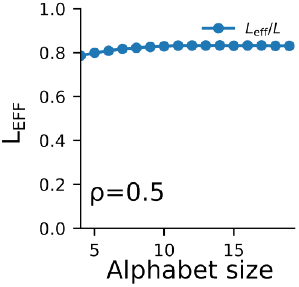
Alphabet size effect on *L*_*eff*_ . Synthetic MSAs of size N=500 and *ρ* = 0.5 but alphabet size *k* is varying from 4 to 20. The *L*_*eff*_ is recorded and shown in blue.

**Figure SI4:**
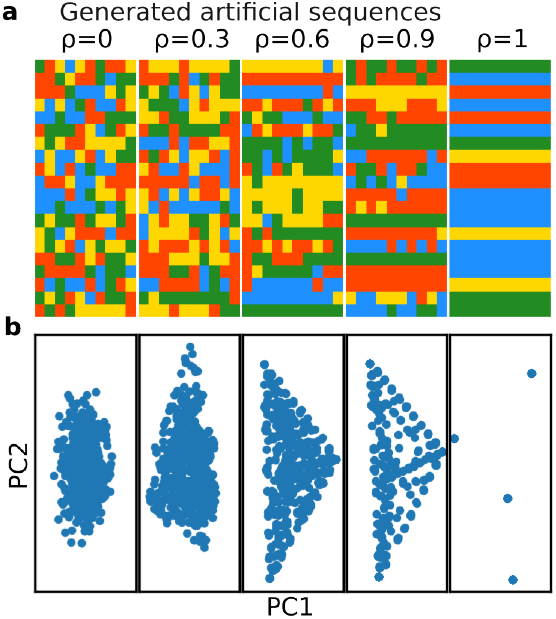
Synthetic MSA with varying correlation strength between positions and symbols. In the first row (a), I show the MSA, where one colour corresponds to a nucleotide and each row is a sequence. In the second row (b), I show the PCA projection of the MSA, showing how they converge to four typical sequences.

**Figure SI5:**
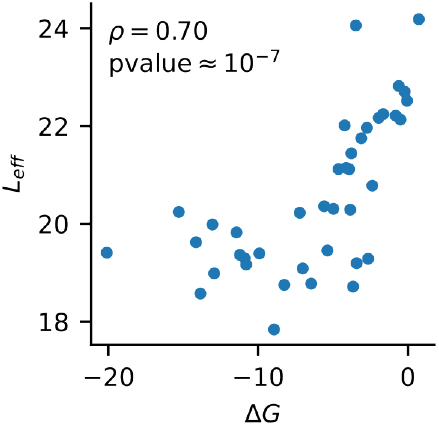
Folding Δ*G* correlates with *L*_*eff*_ . I generated 500 variants using RNAinverse on 40 randomly generated structures. I computed *L*_*eff*_ for each of the 40 datasets together with the average folding stability (computed with ViennaRNA [16]), which is shown as a scatter plot.

**Figure SI6:**
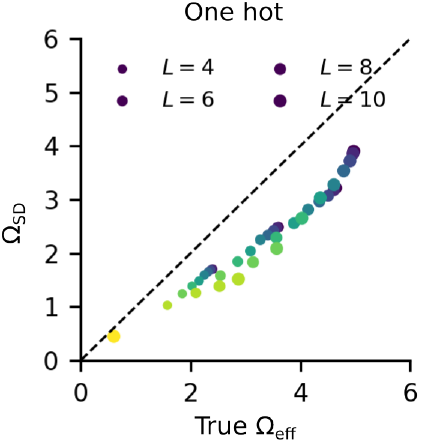
Scatter plot of the estimated effective support size 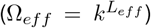 using directly the one hot encoding instead of the Helmert encoding, colored by sequence length *L* (4 to 10). The dashed line represents the identity function (*y* = *x*).

**Figure SI7:**
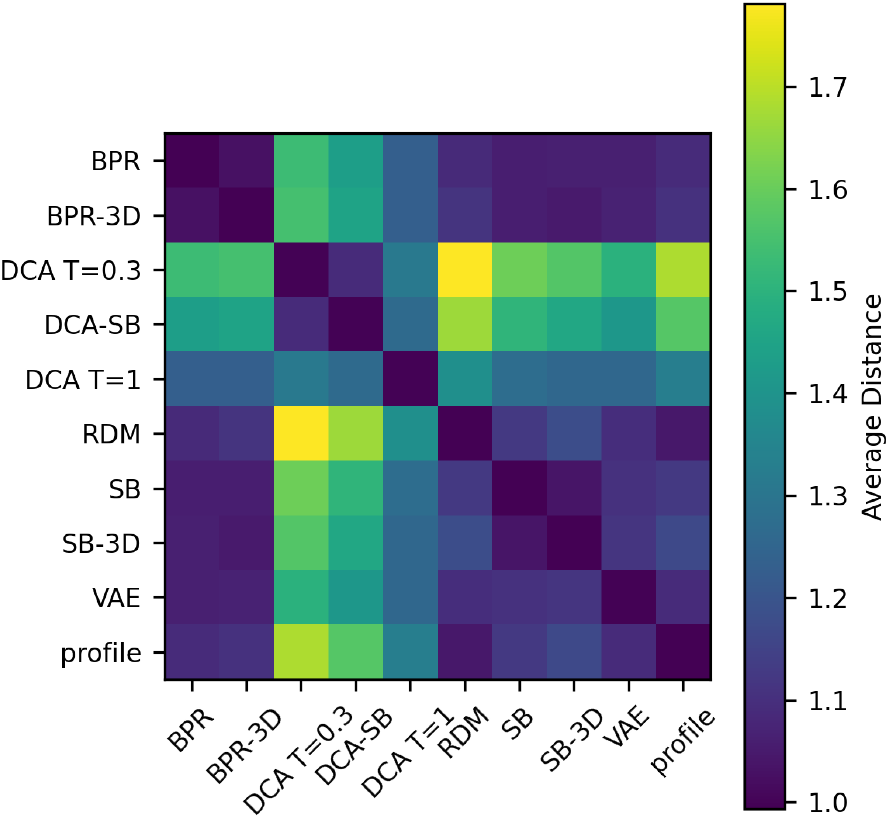
Cross average pairwise distance between generative libraries. Heatmap of the normalized mean Hamming distance computed between RNA sequence libraries produced by each generative model. Sequences are one-hot encoded and distances are averaged over all cross-library pairs. Each cell corresponds to the mean cross distance between library X and library Y, normalized by the geometric mean of their respective within-library average distances. Axes list the generative models, and the color scale indicates the resulting normalized average distance.

